# Feature attention graph neural network for estimating brain age and identifying important neural connections in mouse models of genetic risk for Alzheimer’s disease

**DOI:** 10.1101/2023.12.13.571574

**Authors:** Hae Sol Moon, Ali Mahzarnia, Jacques Stout, Robert J Anderson, Cristian T. Badea, Alexandra Badea

## Abstract

Alzheimer’s disease (AD) remains one of the most extensively researched neurodegenerative disorders due to its widespread prevalence and complex risk factors. Age is a crucial risk factor for AD, which can be estimated by the disparity between physiological age and estimated brain age. To model AD risk more effectively, integrating biological, genetic, and cognitive markers is essential. Here, we utilized mouse models expressing the major APOE human alleles and human nitric oxide synthase 2 to replicate genetic risk for AD and a humanized innate immune response. We estimated brain age employing a multivariate dataset that includes brain connectomes, APOE genotype, subject traits such as age and sex, and behavioral data. Our methodology used Feature Attention Graph Neural Networks (FAGNN) for integrating different data types. Behavioral data were processed with a 2D Convolutional Neural Network (CNN), subject traits with a 1D CNN, brain connectomes through a Graph Neural Network using quadrant attention module. The model yielded a mean absolute error for age prediction of 31.85 days, with a root mean squared error of 41.84 days, outperforming other, reduced models. In addition, FAGNN identified key brain connections involved in the aging process. The highest weights were assigned to the connections between cingulum and corpus callosum, striatum, hippocampus, thalamus, hypothalamus, cerebellum, and piriform cortex. Our study demonstrates the feasibility of predicting brain age in models of aging and genetic risk for AD. To verify the validity of our findings, we compared Fractional Anisotropy (FA) along the tracts of regions with the highest connectivity, the Return-to-Origin Probability (RTOP), Return-to-Plane Probability (RTPP), and Return-to-Axis Probability (RTAP), which showed significant differences between young, middle-aged, and old age groups. Younger mice exhibited higher FA, RTOP, RTAP, and RTPP compared to older groups in the selected connections, suggesting that degradation of white matter tracts plays a critical role in aging and for FAGNN’s selections. Our analysis suggests a potential neuroprotective role of APOE2, relative to APOE3 and APOE4, where APOE2 appears to mitigate age-related changes. Our findings highlighted a complex interplay of genetics and brain aging in the context of AD risk modeling.

## Introduction

Alzheimer’s Disease (AD) is a devastating and irreversible neurodegenerative disorder, affecting millions of people worldwide and posing significant challenges to healthcare systems and society (“2023 Alzheimer’s disease facts and figures,” 2023). The multifactorial etiology of AD, which encompasses genetic, environmental, and lifestyle factors, has made the comprehensive understanding of it complex but essential (Holtzman et al., 2011). Investigating the underlying risk factors associated with AD and their interactions has become a crucial task in neuroscience research (Jack et al., 2018). One of the promising methodologies is the study of brain connectivity through diffusion magnetic resonance imaging (MRI) because of its potential to enhance our understanding of brain networks and their vulnerability, in association with cognition (Le Bihan et al., 2001). Tractography based on diffusion MRI allows construction of structural connectomes (Calabrese et al., 2015; Hagmann et al., 2008; Jones et al., 2013), and modeling properties of the connections between brain regions. Mouse models, given their genetic modifiability, serve as a particularly robust research tool, that allows for high-resolution ex vivo MRI (Badea & Johnson, 2013), elucidating detailed neural circuitry and its potential aberrations in models of AD (Badea et al., 2010; Badea, Stout, et al., 2022; Badea et al., 2019; Daianu et al., 2015; Kerbler et al., 2013; Van der Linden & Hoehn, 2022; Zerbi et al., 2013).

The heterogeneity of lifestyle experience, environmental and genetic traits lead to complex multifaceted neurobiological changes during aging, some of which increase risk for neurodegenerative conditions like AD (Fjell et al., 2014). Age is the most significant risk factor for the development of AD, and brain pathology may be present decades before clinical diagnosis (Sperling et al., 2011). Thus, understanding age-related alterations in the brain is imperative. Neuroimaging based models hold immense potential, not just as diagnostic and prediction tools, but also as markers of disease progression, stratification, and the efficacy of therapeutic interventions (James H. Cole et al., 2017; Franke et al., 2010). The contrast between an individual’s brain age and their chronological age can provide a robust metric to ascertain AD risk, and to locate vulnerable sub-networks, which in turn can suggest targeted early interventions (Gaser et al., 2013; Johnson et al., 2018).

Statistical methods from regression to support vector machines and Gaussian processes have been employed to predict disease conversion, e.g. from MCI to AD (Gaser et al., 2013), and more recently for brain age prediction (Irimia et al., 2015), with a great shift towards deep learning approaches in recent years (Feng et al., 2020), in particular based on single (J. H. Cole et al., 2017), and also multiple imaging protocols (He et al., 2021; Liem et al., 2017). Global and local transformers have been employed to make use of global and local information (He et al., 2022). New GPU architectures, and large public MRI databases have contributed to an accelerated development of image-based models to classify subjects in disease groups, predict age (Antipov et al., 2017; J. H. Cole et al., 2017; Jonsson et al., 2019; Peng et al., 2021; Wang et al., 2019), and segment lesions (Moon et al., 2022), etc. CNNs have been particularly pervasive due to their prowess in handling high-dimensional data like fMRI (Kaufmann et al., 2019), but the literature on structural connectome-based age prediction is still sparse (Bass et al., 2023; Vakli et al., 2018). Graph neural networks (GNN) were found to be well suited for dealing with fMRI-derived connectomes (Ktena et al.; Li et al., 2021) and structural connectivity (Lin et al., 2016), which can benefit from multimodal integration with structural MRI using graph transformers (Cai et al., 2023). Such models may benefit from age-specific stratification, and from integration of external data, allowing one to examine the relationship between cognition and brain imaging derived estimates (Lee, 2023; Millar et al., 2023). Yet, despite these advances, diffusion MRI, with its ability to provide brain structural connectivity, is yet to be fully explored using neural networks in humans, and remains largely unexplored in mice.

Addressing the unique challenges of integrating mouse brain connectomes with other traits for continuous age prediction requires innovation. Recognizing that age is a continuous and not a discrete variable necessitates a shift from classification to prediction frameworks. Historically the focus has been on brain regions, which make up the nodes in a connectome, but understanding AD’s progression should address changes in the edges connecting such nodes (Hagmann et al., 2008). Many of the previous studies have focused on capturing salient nodes from functional connectomes. There is a need to map important edges reliably from structural connectomes. In order to discern the important connections, there is a need to minimize connectomic noise and mitigate potential bias and overfitting. Further, the small sample sizes and data heterogeneity from genetic and environmental factors compound these challenges (Bettio et al., 2017; Liu et al., 2015). Mouse models, with their uniform genetic backgrounds and controlled environments, allow smaller sample sizes to inform on the effect of genetic risk on the connectomes. However, our mouse studies feature relatively larger image sizes, accompanied by adjacency matrices of 332 x 332 connections (Anderson et al., 2019; Calabrese et al., 2015).

To address our objective of brain age prediction in a mouse population, we propose a Feature Attention Graph Neural Network (FAGNN) model. The proposed FAGNN’s architecture utilizes distinct sub-networks tailored to process each data type, encompassing connectomes, risk factors and behavioral data, thereby ensuring a more robust and comprehensive analysis (Raj et al., 2010; Smith & Topin, 2019). Drawing inspiration from the multi-head attention mechanisms of transformer models (Vaswani et al., 2017), we developed a quadrant attention module (QAM) to capture the interplay of brain connections more efficiently by processing each quadrant of the connectome separately as feature dimensions. This approach enables the QAM within FAGNN to efficiently capture and highlight task-specific brain connections, leveraging unique data from each connectome quadrant for comprehensive analysis.

In our study, we aimed to validate the network’s identification of important structural connections by examining key tractography features that influence the structural connectome. We initially utilized diffusion weighted imaging (DWI) metrics, specifically fractional anisotropy (FA), to assess the directional diffusion of water molecules within brain tissues. FA provides valuable insights into the microstructural integrity of white matter tracts. However, it is based on the assumption of Gaussian water diffusion, which may not adequately represent the complexity of brain tissue microstructures. To address the non-Gaussian nature of water diffusion in the brain, especially in areas with complex fiber arrangements, we included mean apparent propagator (MAP) metrics. MAP metrics, derived from models such as the simple harmonic oscillator-based reconstruction and estimation (SHORE), offer a more accurate representation of the diffusion process in brain tissues. These metrics include return-to-origin probability (RTOP), return-to-axis probability (RTAP), and return-to-plane probability (RTPP), which correspond to various aspects of pore structure as described in pore theory (Callaghan et al., 1992). Specifically, RTOP is associated with the average volume of pores, RTAP to their cross-sectional area and RTPP to their length as detailed in earlier studies, meaning higher values of these metrics reflect a more restricted diffusion in neural tissues (Bogusz et al., 2022; Mitra et al., 1995; Özarslan et al., 2013). These quantitative indices offer valuable information of tissue heterogeneity (Brusini et al., 2018), ischemic stroke (Brusini et al., 2016; Galazzo et al., 2018), multiple sclerosis (Brusini et al., 2021), cognitive impairment detection (Menon et al., 2020; Pitteri et al., 2021), and aging and dementia (Bouhrara et al., 2023).

The primary objective of this study was to explore the potential of advanced neural network architectures, specifically the FAGNN, for predicting brain age, as age is the most significant risk factor for AD. By utilizing a deep learning-based integrative modeling approach that incorporates multivariate datasets including brain connectomes, traits such as sex and genotypes, and cognitive metrics, we aim to reveal the interactions between such factors and the aging process. We sought to identify specific brain connections that play a significant role in age prediction, offering deeper insights into possible association with the progression of brain aging. Our approach can serve to find early biomarkers related to aging, to facilitate the design and testing of future preventative, and therapeutic interventions.

## Methods

### Animal Models

In this study, we examined the effects of five contributing factors to Alzheimer’s disease risk on brain connectomes: APOE genotype, presence of human NOS2 gene, diet, age, and sex. We used animal models with the mouse APOE gene replaced with the three major human APOE variants. These mice were homozygous for the APOE2, APOE3 and APOE4 alleles. To better reflect the human innate immune system, we included mice with a human NOS2 (HN) gene (Colton et al., 2008). To address sex as a biological variable, and increased risk for AD, our sample selection included male and female mice. In our study of age-related risk factors, we included a total of 170 mice separated into two cohorts: a group of 66 mice aged 12 to 15 months to represent middle age, and a group of 104 mice aged 15 to 20 months to represent older age. To introduce an environmental risk factor, we subjected mice to either a standard or high-fat diet. The animal groups were distributed across APOE genotypes (58 APOE2, 56 APOE3 and 56 APOE4 homozygous mice), sex (82 males and 88 females), HN (95 mice with no HN gene and 75 with HN gene, and diet (113 control diet and 57 high-fat diet).

### Image Acquisition and Connectomes

To derive structural connectomes, we used high field, 9.4T MRI to image fixed brain specimens, using a previously published protocol (Badea, Li, et al., 2022; Winter et al., 2023). To prepare the brain specimens for imaging, mice were sacrificed through a transcardiac perfusion fixation with 10% formalin and 10% (50 mMol) gadoteridol (ProHance, Bracco), then the mouse brains were trimmed and immersed in 0.05% (2.5 mMol) ProHance with PBS (0.01 molar, pH 7.4). A compressed sensing 3D SE sequence was used for DWI, with the following parameters: TR/TE of 100 ms/14.2 ms, and BW of 62.5 kHz. Diffusion pulses were applied in 46 directions with 2 diffusion shells (23 at 2,000 and 23 at 4,000 s/mm^2^) and we used 5 non-diffusion-weighted (b0) images. The diffusion pulse has a maximum amplitude of 130.57 Gauss/cm, a duration of 4 ms, and a separation of 6 ms, with eight-fold compressed sensing acceleration (Anderson et al., 2019; Wang et al., 2018b). A matrix size of 420 x 256 x 256 and FOV of 18.9 mm x 11.5 mm x 11.5 mm produces images with 45 μm isotropic resolution. We used MRtrix3 (Tournier et al., 2019) to create tractography and connectomes based on the acquired diffusion MRI images. Approximately 10 million tracts were generated, from which a subset of 2 million tracts was selected for each subject, based on a FA cutoff of 0.1 and a maximum length of 1000 mm. We calculated the microstructural parameters based on diffusion using MRtrix3 (Tournier et al., 2019), and RTOP, RTAP and RTPP using Dipy (Garyfallidis et al., 2014).

### Behavioral Metrics

Mice with genetic and environmental risk for AD can exhibit cognitive decline, including learning and memory deficits, which are characteristic of aging and AD. To quantitatively assess behavioral differences, we employed the Morris Water Maze (MWM) test. Mice were introduced to a pool filled with room temperature water, rendered opaque by the addition of nontoxic white paint. The pool was divided into four quadrants (SW, NE, NW, SE), a thigmotaxis area, and a hidden platform located in the SW quadrant’s center. During training trials, the mice were released from each quadrant for four trials per day for 5 consecutive days and had to find the hidden, submerged platform using external visual cues, swimming for a maximum of 60s. We measured the time and distance to platform and quadrant-specific measures to evaluate spatial learning and memory. Two probe tests were conducted: at one-hour after the final training trial (day 5), and another one at 72 hours (day 8) to evaluate memory retention for the platform location. Metrics included swim distance, swim velocity, time to locate platforms, time and distance in each quadrant, number of entries to probe location and winding numbers representing rotational movement during swimming (Badea, Li, et al., 2022).

### Feature Attention Graph Neural Network (FAGNN)

Binary risk factors such as sex, diet and the presence of the HN gene were normalized to values of 0 or 1. The genotype data of APOE alleles (2, 3, and 4) was scaled to 0, 0.5, and 1 respectively. We ranked normalized connectome data, representing the connections between different brain regions. Equal numbers of connections were assigned the same rank, which was then divided by the maximum rank, and scaled between 0 and 1. Behavioral data was also normalized, with time-based metrics divided by 60, which is the maximum time duration (in seconds) of each MWM trial, and other metrics divided by their corresponding maximum values to bring them to a similar scale. Thus, the entire data used in the network was scaled between 0 and 1.

Our integrative modeling approach for age prediction (**Figure 1**) includes multiple specialized sub-networks for each input modality. The behavioral data is fed through a 2D Convolutional Neural Network (CNN). The arrangement of the behavioral data ensures that similar metrics placed adjacently in columns and trials progressing from earlier to later are represented in a spatially contiguous manner, which aids the 2D CNN in effectively extracting spatial features and patterns that are relevant to the progression of the behavioral data. Risk factor data based on subject traits (age, sex, genotype), being one-dimensional, are passed through a 1D CNN. This is effective for capturing patterns across the 1D data especially given that we later combine these output tensors with those of 2D CNN for tensor homogeneity. This captures cognitive markers associated with disease biomarkers. Connectome data is first passed through a quadrant attention module (QAM), which is responsible for assigning importance scores to different brain connections. This helps in highlighting the most important connections in the brain with respect to AD risk factors. After the QAM, the connectome graph data is multiplied elementwise with the edge scores, then processed by a Graph Neural Network (GNN), which incorporates Graph Convolutional Networks (GCNs) and top-K pooling layers (Gao & Ji, 2019). The general structure of the GNN sub-network is based on BrainGNN (Li et al., 2021) which was originally designed for processing fMRI connectomes for classification. We modified the model by including a mean-squared error loss function to predict age, which is a continuous variable. The top-K pooling layers in the GNN help in selecting the top K percent of nodes in each graph layer, based on the nodes’ importance for decreasing noise, and dimensionality reduction. The output tensors from each sub-network were concatenated and passed through a fully connected (FC) block.

**Figure 1.**
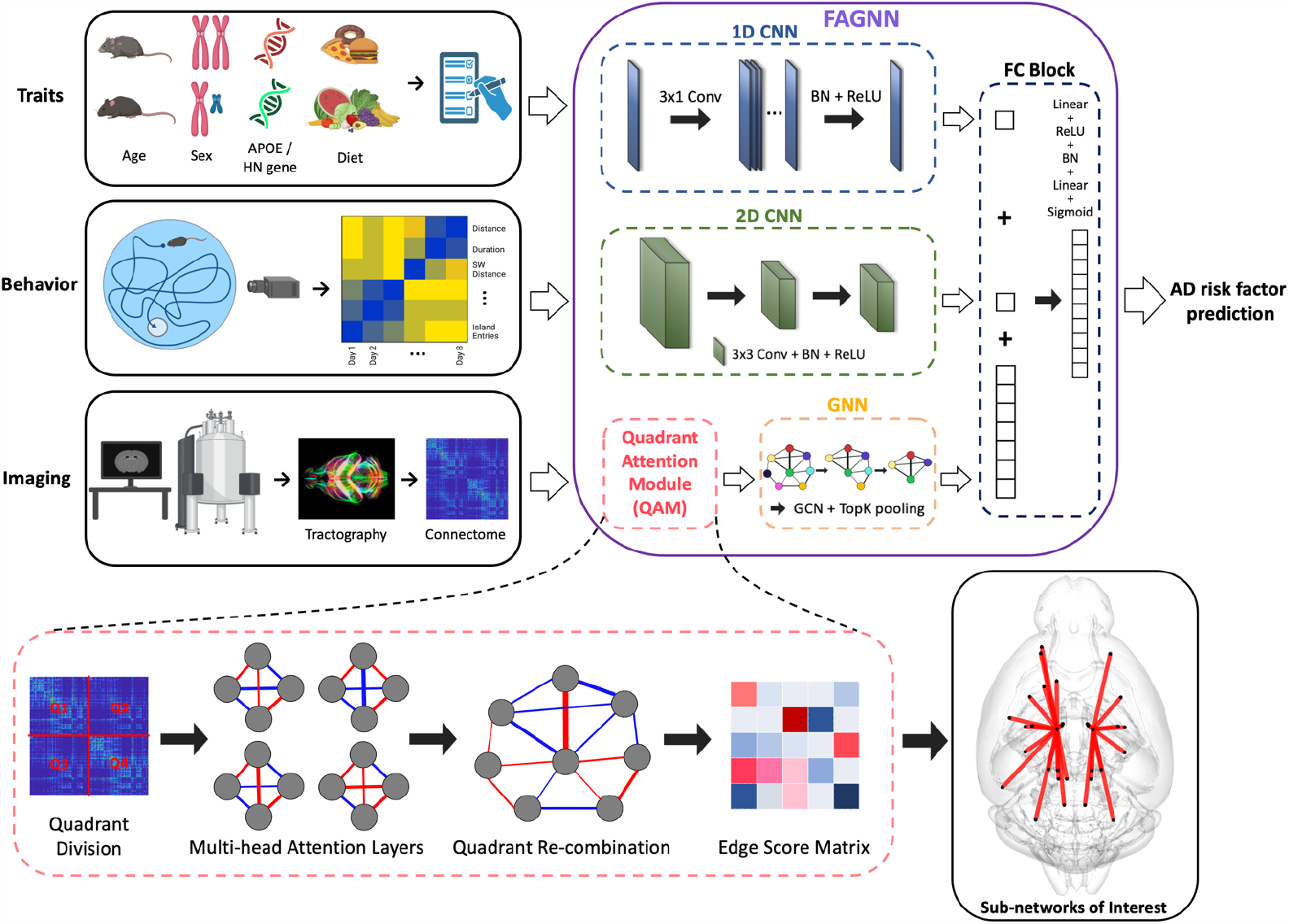
Overview of FAGNN method integrating AD risk factor traits, behavioral metrics and brain connectome. Diffusion MRI-derived connectomes undergo quadrant attention module for edge scoring prior to GNN analysis. AD risk traits and behavioral metrics are processed by 1D and 2D CNNs, respectively. The network outputs predicted brain age and we determine the difference between brain age and chronological age as a risk factor.

In our QAM, we leverage the structure of brain connectome data, which is organized in a matrix form of size *n* × *n*, representing the interconnectivity between *n* different regions of the brain. The QAM addresses the high dimensionality and potential symmetry within the connectome by dividing the connectome matrix into four quadrants and applying a distinct multi-head attention (MHA) mechanism to each quadrant.

Given an input connectome data tensor *X* ∈ ℝ ^*n* × *n*^, we partition it into four quadrants:

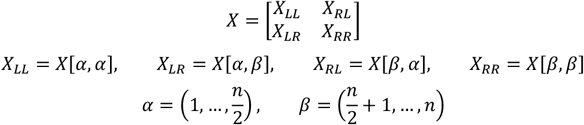

Each quadrant *X*_*pq*_, with p, q is then processed through MHA separately. The MHA operation is formulated as follows (Vaswani et al., 2017):

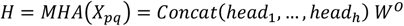

Here, each attention head *head*_*j*_ for *j* = 1, …, *h* is computed using the scaled dot product attention mechanism:

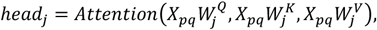

where 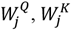 and 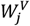 are the weight matrices for the query (Q), key (K) and value (V) respectively for *head*_*j*_, and *W*^*O*^ is the output weight matrix that combines the heads. The attention function is defined as:

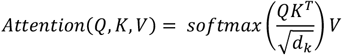

where *d*_*k*_ is the dimension of each key. In the QAM, this scaled dot-product attention mechanism is applied independently to each quadrant of the connectome. The four quadrants correspond to the brain’s hemispheric divisions: left-to-left (LL), right-to-right (RR), left-to-right (LR) and right-to-left (RL) hemispheric connections. This quadrant-specific processing aligns with the brain’s method of regional communication through inter- and intra-hemispheric connections. Thus, the QAM in our model narrows the focus to distinct quadrants for a more targeted analysis of age-related neural connections. By utilizing MHA layers for each quadrant, QAM facilitates a detailed yet efficient edge score calculation of the connectome. This approach not only aids in managing larger datasets but also optimizes computational resources, contributing to the adaptability and practicality of our model in various research settings.

Then, we propose a parallel sub-network schematics of GNN and CNN layers:

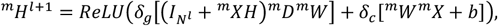

where *m* is the sub-network specification, *l* is the current level of layer, *H* is the edge scores from the quadrant attention module, *δ*_*g*_ is the delta function for GNN layers, *δ*_*c*_ is the delta function for CNN layers, 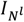 is the identity matrix with dimension of the number of nodes at layer *l, X*_*m*_ is the connectivity matrix, *D* is the diagonal matrix of the graph data, and *W*_*m*_ is the learnable model parameters of sub-network *m*.

After the QAM, the FAGNN is employed to process the connectomic graph data. FAGNN integrates GCNs and CNNs to perform non-linear transformations, computing the subsequent hidden state *H* ^*l* +1^ based on the output from the quadrant attention module *H*. This enables FAGNN to efficiently represent the graph structure of connectome, while accounting for the attributes of nodes and edges within the graph. Moreover, given that the dataset encompasses mice exhibiting multiple risk factors, the inclusion of risk factor data in the prediction helps the model to discern and account for the potential influence of various risk factors and their intricate interplay on brain connectivity, ensuring a less biased representation. Additionally, the incorporation of behavioral data is necessary, as it represents cognitive functions, enabling the model to account for potential correlations with brain connectivity patterns and AD risk factors. By integrating these elements, FAGNN contributes to a more holistic comprehension of the interplay between brain connectivity, cognitive performance, and AD risk factors.

We used an Adam Optimizer with multi-learning rates for the different sub-networks:

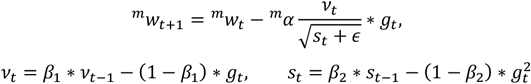

where ^*m*^*w*_*t*_ represents the parameters of the network being optimized for sub-network *m* at time *t*, ^*m*^*α* is the learning rate for sub-network *m, g*_*t*_ is the gradient at time *t* along w, *v*_*t*_ is the exponential average of gradients along *w, s*_*t*_ is the exponential average of squares of gradients along *w*, and *β*_1_ and, *β*_2_ are the hyperparameters; *ϵ*, a small non-zero value to ensure there is no division by zero, is set at 1e-10.

To train the model efficiently, we used different learning rates for different sub-networks, which is helpful considering the different types of data being processed by each sub-network. The optimizer computes the moving averages of gradients and their squares and uses them to adaptively adjust the learning rates during training. This helps in achieving faster convergence during training and can lead to better performance of the model. Each sub-network was assigned its unique learning rate depending on the performance. The full model integrates sub-networks including QAM, GNN, 1D CNN, 2D CNN, and FC block. The respective learning rates used were 0.001, 0.001, 0.002, 0.002 and 0.003 with 100 epochs.

In our model evaluation, we utilized a 5-fold cross-validation approach, where the dataset was divided into five distinct folds for comprehensive training and testing. This methodology allowed us to compare the performance of the FAGNN model against several variations: Comp 1 utilized connectome data processed via a modified BrainGNN, Comp 2 combined the QAM with the connectome, Comp 3 added behavioral data to the QAM and connectome, Comp 4 integrated trait data with the QAM and connectome, and the full FAGNN model included the QAM, connectome, trait data, and behavioral data. The accuracy of these models was quantified by calculating the mean absolute error (MAE) and root-mean-squared error (RMSE), with each model run repeated 10 times to ensure consistency in the results. For our statistical analyses, we employed linear models within the R programming framework, considering a p-value less than 0.05 to be statistically significant.

For a tract-based analysis of FA and MAP parameters, we divided the animals into three age groups, young (12-14 months), middle (14-16 months), and old (>16-20 months). This categorization imposes granularity, or stratification, to our assessment of microstructural changes based on DTI and MAP metrics across aging stages. Linear mixed models were used for tract-based comparisons. Post hoc pairwise comparisons were conducted using Dunn’s test to calculate the significance of difference of each group separately after a non-parametric Kruskal-Wallis test. Considering the multiple comparisons being made, p-values from the Dunn’s test were adjusted using the Holm method to control the familywise error rate. We trained the models using a Lambda server equipped with four RTX 8000 cards, each with 48 GB of memory. The results were visualized using DSI studio (http://dsi-studio.labsolver.org) and brainconn2 (Mahzarnia et al., 2023).

## Results

To help estimate risk for AD based on the discrepancy between chronological and brain age, we built a model for brain age prediction with a dataset comprised of 170 mice. Our strategy used 5-fold cross testing, with each fold including 136 mice used for training, and 34 mice used for testing; and each fold was trained for 10 runs. Along with age prediction, FAGNN results included important edges and nodes. We validated these connections through a comparison of diffusion-parameters using FA and MAP-based metrics associated with tractography, to test whether microstructural parameters support the connectivity changes during aging.

### Model performance in age prediction

The Feature Attention Graph Neural Network (FAGNN) was assessed for its capability to estimate brain age from structural connectome data. The model’s performance was evaluated by its MAE of 31.85 days and RMSE of 41.84 days, as shown in **Table 1**. The FAGNN model outperformed a modified BrainGNN and other configurations of specific data and corresponding sub-network inclusion. The second-best model included the quadrant attention module with connectome risk factors (Comp 4), followed by a similar model which included behavior data instead of the risk factors (Comp 3). The next ranked model included only the quadrant attention module with connectome (Comp 2), and the lowest ranked model only included connectome data through a modified BrainGNN (Comp 1). These measures of accuracy indicated the FAGNN full model’s proficiency in estimating brain age, which is a conceptual age derived from brain connectome patterns, and thus may differ from chronological age. This estimation forms the basis for subsequent analyses of brain aging and the impact of various risk factors.

**Table 1.**
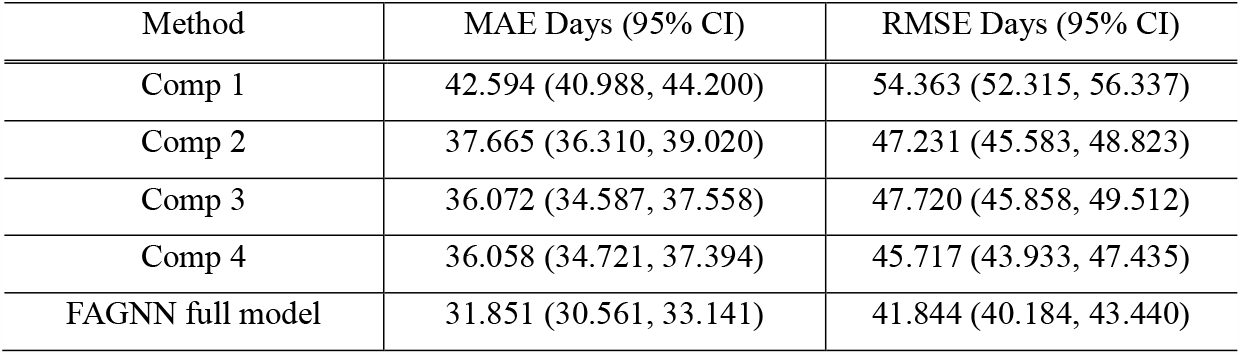
Age prediction performance of various combinations of GNN and FAGNN. The testing metrics include MAE and RMSE. Comp1: connectome (modified BrainGNN); Comp2: QAM + connectome; Comp3: QAM + connectome + behavior; Comp4: QAM + connectome + risk factors. FAGNN full model included QAM + connectome + risk factors + behavior.

### Impact of risk factors on brain age

We estimated the Mean Estimation Difference (MED) between the predicted brain age and the subjects’ actual chronological age. This difference provides insight into the neuroprotective or neurodegenerative influences exerted by genetic and environmental factors. A lower MED indicates a younger brain age relative to chronological age, suggesting potential resilience against aging-related changes. **Figure 2** depicts how different AD risk factors were associated with variations in MED, emphasizing their potential impact on the aging brain. With respect to the effect of genotypes, APOE2 had lowest MED of the three genotypes followed by APOE4 and APOE3. Sex effects indicated that females had lower MED values than males. The presence of the HN gene and high fat diet yielded higher MED than the wildtype and control diet mice.

**Figure 2.**
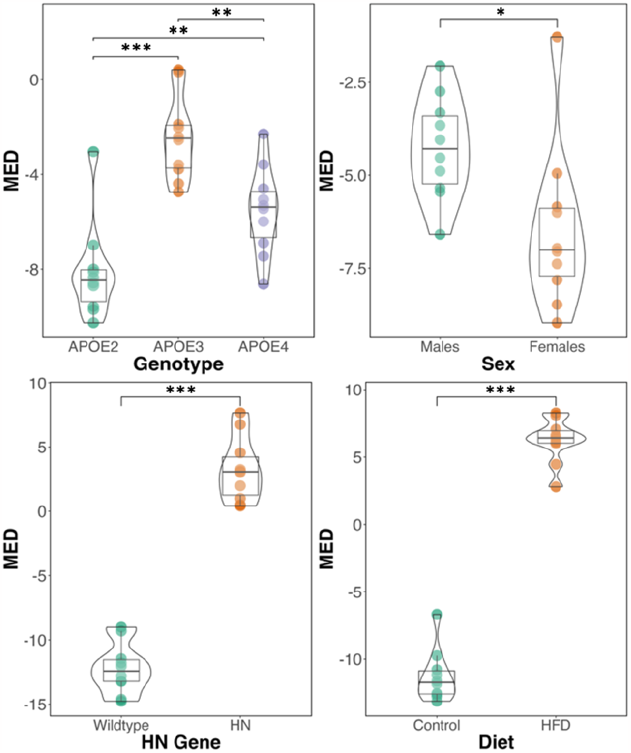
Mean estimation difference (MED) of brain age for the risk factors used in the model. The risk factors include APOE genotype, sex, presence of HN gene and diet. (*** p<0.001, **p<0.01, *p<0.05).

### Connectomic and tractography analysis across age groups

The FAGNN model identified the top 30 brain connections that were most influential in age prediction (**Figure 3**). These pointed to a central role for the cingulum and its connections. Other nodes included the piriform cortex, frontal association cortex, primary and secondary motor cortex, striatum, amygdala, hippocampus, thalamus, midbrain reticular nucleus, pontine reticular nucleus, caudomedial entorhinal cortex, cerebellar cortex, and brain stem. Thus, age affected widespread brain networks, including regions with known roles in memory and motor function, and we identified a central role for cingulum.

**Figure 3.**
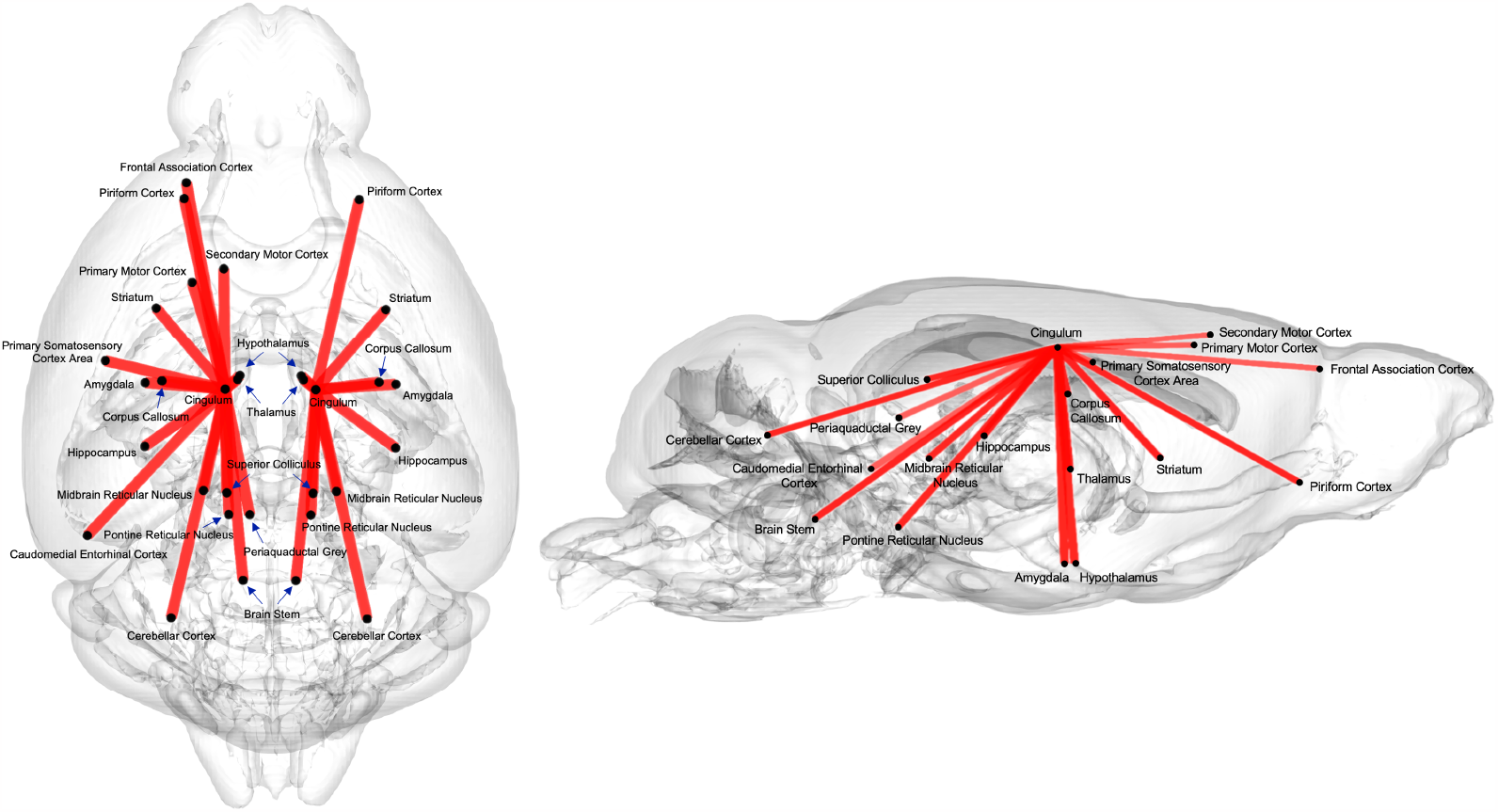
The top 30 connections with highest edge scores from continuous age prediction given by FAGNN. These connections were associated the most with age variation. The regions are connected to either the left or right cingulum forming two separate but mostly symmetric sub-networks.

To validate the output connections from the QAM, we performed a tractography analysis examining the FA and RTOP values across the top 12 tracts identified by the model (**Figure 4**). The top 12 tracts include both left and right intra-hemispheric connection between striatum, corpus callosum, hippocampus, brain stem, piriform cortex and thalamus to cingulum. We visualized these tracts to show how they are spatially arranged, and the variability in FA and RTOP.

**Figure 4.**
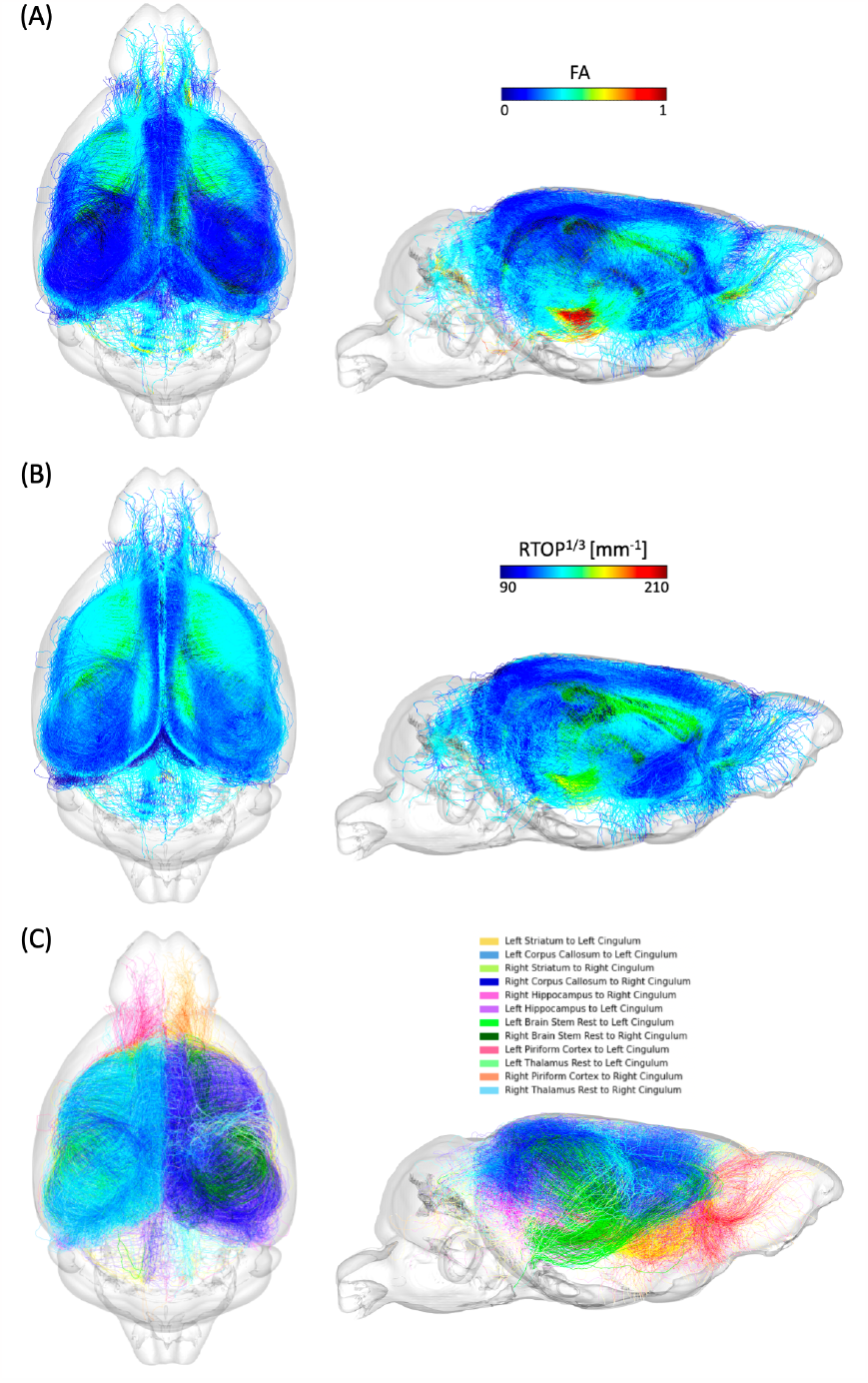
Visualization of the top 12 connections that contributed most to age prediction based on the quadrant attention module. (A) FA values along the corresponding tracts. (B) RTOP values along the corresponding tracts. (C) Illustration of the tractography with distinct colors for each connection.

Tract-specific analyses were conducted across the three age groups (young, middle-aged, and old age) for the top 6 tracts identified by FAGNN, which included cingulum to striatum, corpus callosum, and hippocampus for both left and right hemispheres. These analyses revealed how the brain’s structural integrity along tracts varies with aging, i.e., FA and RTOP decreased with age. The results, visualized in **Figures 5** and **Figure 6**, support the FAGNN model’s selections and offer insight into the microstructural properties of the tracts that may underpin age-related cognitive changes. Specifically, the FA analysis is shown in **Figure 5** and the RTOP analysis is shown in **Figure 6**.

**Figure 5.**
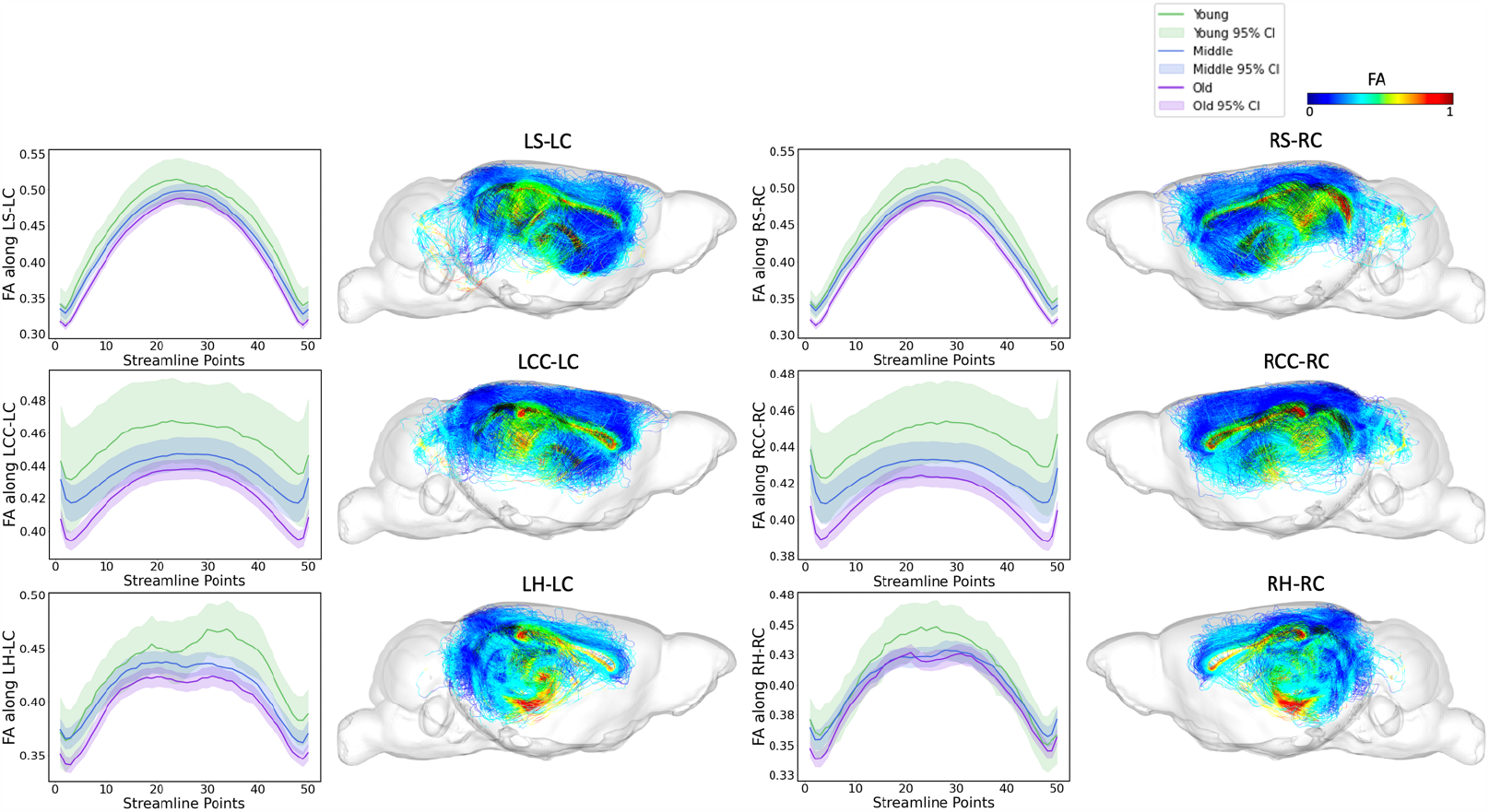
FA along tract profiles for the top 6 edges identified by FAGNN. The first row shows striatum-cingulum tract, second row shows corpus callosum-cingulum tract and last row shows hippocampus-cingulum tract. First column shows FA profile of left-left connection, second column shows the corresponding tractography with FA values. Third column shows FA profile of right-right connection, and the last column shows the corresponding tractography with FA values. The FA values along each tract for age groups were significantly different with p<0.001.

**Figure 6.**
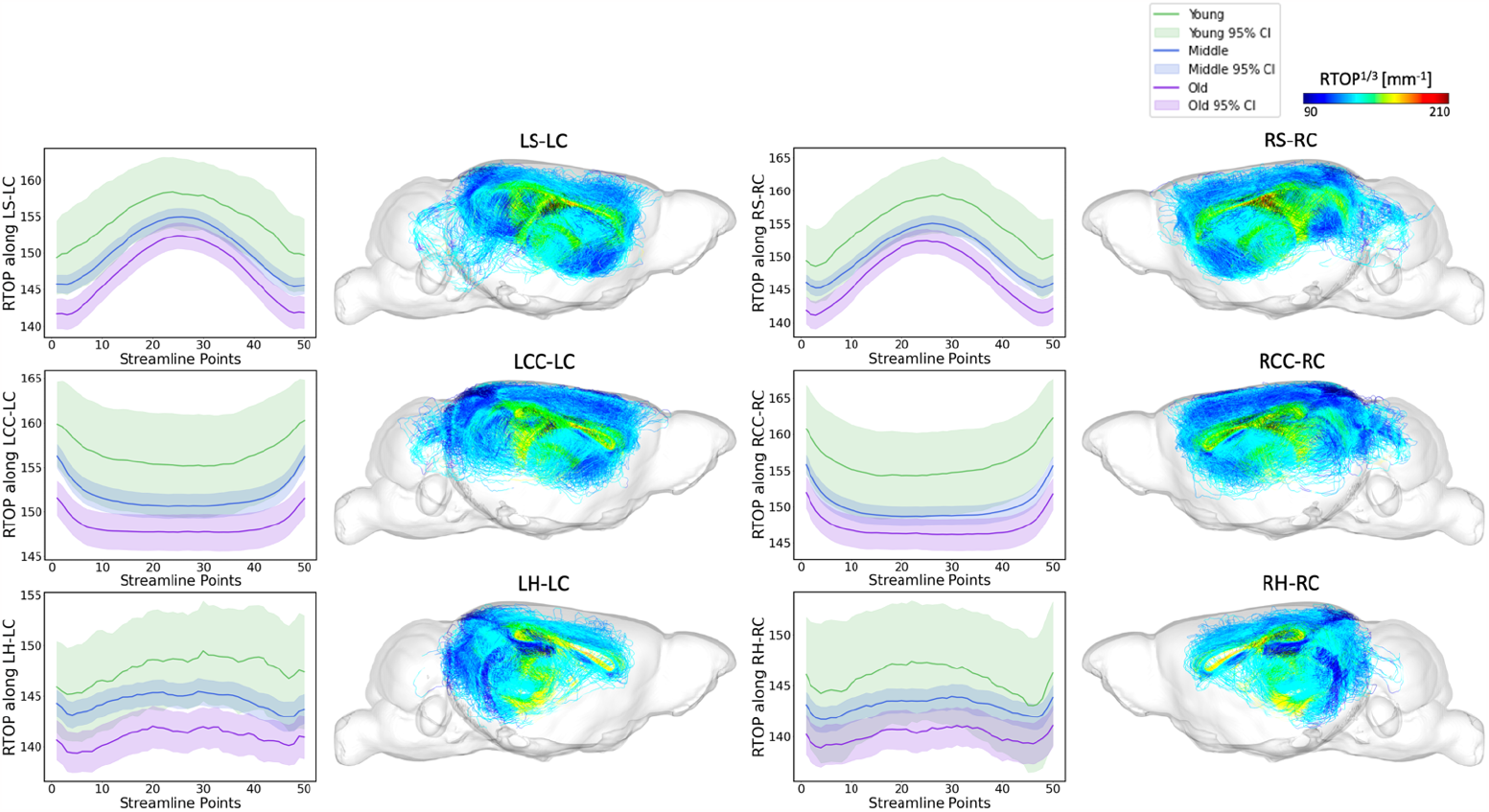
RTOP along tract profiles for the top 6 edges identified by FAGNN. The first row shows striatum-cingulum tract, second row shows corpus callosum-cingulum tract and last row shows hippocampus-cingulum tract. First column shows RTOP profile of left-left connection, second column shows the corresponding tractography with RTOP values. Third column shows RTOP profile of right-right connection, and the last column shows the corresponding tractography with RTOP values. The RTOP values along each tract for age groups were all significantly different with p<0.001. **Figure 7**. and **Figure 8** extend these analyses to include RTAP and RTPP values, further characterizing the microstructural details of these tracts and supporting the biological relevance of the connectomic features identified by the FAGNN model as markers of brain aging.

The overarching pattern emerging from our analyses is that young mice have higher white matter microstructural metric values including FA along the FAGNN-identified connections, middle-aged mice have middle values and older mice have lower values. These metrics decreased with increasing age. The FA and MAP related values along each tract for age groups were all found significantly different with p<0.001. RTOP, RTAP and RTPP respect the same pattern as observed for FA variations with age.

## Discussion

The FAGNN introduced in this study represents a novel approach for predicting brain age from multivariate data (connectome, behavior and traits) from mouse models of Alzheimer’s disease risk. The model yielded predictive accuracy with a mean absolute error of 31.85 days and a root mean squared error of 41.84 days, and outperformed other networks, indicating its reliable degree of precision in estimating brain age. This suggests FAGNN’s potential to significantly enhance our understanding of the relations between brain connectivity and aging. To establish the efficacy of FAGNN, we conducted a comparative analysis with a modified BrainGNN (Li et al., 2021), which was originally designed for functional connectomes and for classification tasks, and that we adapted for structural connectomes and for continuous variable prediction tasks, i.e. age.

We evaluated various configurations of our model that included distinct sub-networks to discern which combination and which subsets of the input data most effectively contribute to the accuracy of age prediction. We found that our full model, which integrates connectomes processed within a quadrant attention module, AD risk factor data, and behavioral metrics outperformed all other models we have tested. The better performance of the comprehensive model emphasizes the importance of a multi-dimensional and multi-variate approach integrating diverse datasets for enhancing the model’s predictive accuracy.

Chronological age inherently influences a multitude of factors in mice, extending beyond those related to brain-related traits (De Magalhães et al., 2005; Kovacs et al., 2009). Thus, an accurate prediction of chronological age would necessitate the inclusion of many variables outside the scope of our dataset, which is primarily focused on factors pertinent to brain structural connectivity as a marker of successful or pathological aging. This focus has led to the recognition of a disparity between the chronological age and the brain age predicted by our model, which is estimated based on cognition, AD-related traits and brain connectivity. We thus estimate the discrepancy between chronological and brain age as a marker of risk (Anatürk et al., 2021; Franke & Gaser, 2019). We have used the MED between the chronological brain age and brain age, as estimated by the model (James H. Cole et al., 2017; Grosenick et al., 2013; Le et al., 2018; Liang et al., 2019). A lower MED could suggest that the brain’s connectomic structure with corresponding cognitive traits and AD risk factors are younger than its chronological equivalent, potentially indicating a protective mechanism against age-related changes. The variations in MED observed in our study highlighted the influence of genetic and environmental factors on the aging process of the brain. Mice with an APOE2 genotype had the smallest MED among all of the APOE genotypes, suggesting a possible neuroprotective mechanism against aging, and in agreement with the theory that an APOE2 genotype may confer resilience against brain aging and AD (Moon et al., 2023; Suri et al., 2013). Surprisingly, the MEDs of APOE3 and APOE4 had the opposite trend than expected, which may result from the age heterogeneity and from other AD risk factors in our sample. We found that innate immunity and diet induced significant increases in MED compared to control counterparts. Our findings are consistent with research linking innate immunity and lifestyle factors such as diet, to changes in neuroprotection and resilience against brain aging (Jurgens & Johnson, 2012). In particular, a high-fat diet confers vulnerability to the brain during aging (Uranga et al., 2010). A larger sample would be needed to address APOE genotype specific changes due to immunity and diet.

Our model identified the brain connections significantly contributing to age prediction, potentially mapping sub-networks vulnerable to aging and increased AD risk. The top 30 brain connections, determined by the quadrant attention module of the FAGNN model, were illustrated in **Figure 4.A**. These connections involve the piriform cortex, striatum, hippocampus, amygdala, thalamus, and hypothalamus, all converging on the cingulum. This selection aligns with previous studies highlighting the importance of these regions in aging (Bartsch & Wulff, 2015; Catheline et al., 2010; Fama & Sullivan, 2015; Shen et al., 2013; Swaab et al., 1992; Umegaki et al., 2008). Notably, these connections predominantly involved the cingulum, a white matter tract involved in a wide range of motivational and emotional behaviors, and importantly in spatial working memory (Schmahmann et al., 2007), as tested in the MWM portion of our study. The cingulum, running anterior to posterior, is considered part of the limbic circuit, and is highly integrated with other white matter tracts (Zerbi et al., 2013). It connects numerous regions and partakes in numerous functions related to memory, problem solving, attention, sensory and regularization of heart rate and blood pressure (Gunbey et al., 2014). There have been previous studies that related cingulum to aging. A study found that sensory and motor functions, which are associated with the cingulate, decline with aging and are reflected by changes in the integrity of the white matter tracts (Madden et al., 2012). The top 30 connections linked either the left or right cingulum to other regions, which supports the important role of the cingulum and its sub-networks in aging, and in relation to AD risk (Bennett et al., 2015; Gold et al., 2012). Our results suggest that additional smaller tracts connecting various brain regions to the cingulum play a significant role as well. Previous studies have shown anterior-posterior white matter tracts demyelination with aging (Madden et al., 2012; Sibilia et al., 2017; Sullivan & Pfefferbaum, 2006). These previous studies motivate further studies of white matter integrity and myelination with aging, expanding from tracts such as cingulum to its connections, as identified by the FAGNN model.

The model assigned edge scores to connections based on prediction accuracy, and connections with higher edge scores more significantly influenced the network’s age prediction decision. These selections are complex and not immediately intuitive, which further prompted investigation into the basis of the network’s choices. We aimed to validate the network’s findings by examining tractography features impacting the structural connectome, utilizing diffusion and MAP metrics that relate to the white matter integrity of the tractography in the brain (Avram et al., 2016; Kochunov et al., 2012; Pitteri et al., 2021; Zucchelli et al., 2016). The tracts with FA and RTOP values are depicted in **Figures 4.B** and **C**. Previous studies have found decreases in diffusion metrics--namely FA values in DTI--in the cingulum decrease with aging (Catheline et al., 2010; Jang et al., 2016; Sibilia et al., 2017). Also, cognitive control in older adults is sensitive to variations in cingulum microstructure for subjects with MCI (Metzler-Baddeley et al., 2012). In mice, there was a decrease in myelinated fiber length density and FA in the cingulum bundle with increasing age, regardless of genotype, hemisphere or sex (Segal et al., 2010). In our investigation, we have expanded the study of white matter integrity using MAP properties including RTOP, RTAP and RTPP. Although there has been a stress on the importance of cingulum in previous studies, there is sparse information on the properties of its connections to other regions, especially in mice. We further expanded the information about the properties of the cingulum connections to specific regions such as striatum, corpus callosum and hippocampus, which was made possible due to high resolution ex-vivo mice studies and advances in accelerated acquisitions benefitting from compressed sensing (Anderson et al., 2018; Wang et al., 2018a), and sensitive to diffusion properties (Anderson et al., 2020), which allowed a more detailed representation of tracts and increased sensitivity for detecting differences in connectivity using deep learning models.

We concentrated on the top six connections differentiated by the model in terms of aging, examining the tract profiles of streamlines within these specific connections, including striatum to cingulum, corpus callosum to cingulum, and hippocampus to cingulum, for both left-to-left and right-to-right hemispheric connections. This focus was to uncover the underlying information of the connectivity and validate the model’s selections. The tract profiles were examined to discern metric values along the tracts, differentiating the white matter integrity and characteristics across the age groups in our dataset, categorized as young, middle-aged, and old mice with age cutoff at 14 and 16 months. In **Figure 5**, we observe a significant decrease in FA values in an age-related sequence, with the highest values in young individuals, decreasing through middle-aged individuals, and reaching the lowest values in old individuals. This aligns with expectations and shows results consistent with previous research on the cingulum (Catheline et al., 2010; Jang et al., 2016; Segal et al., 2010; Sibilia et al., 2017). **Figure 6** displays a similar trend in RTOP values, showing high significance. **Figures 7** and **8** present RTAP and RTPP values along the tracts respectively. The general trend for FA, RTOP, RTAP, and RTPP values across age groups was consistent: young subjects exhibited the highest values, middle-aged subjects had intermediate values, and old subjects showed the lowest values. Notable changes in FA, RTOP, RTAP, and RTPP values within these tracts denote significant variations in the microstructural integrity of the brain tracts. A decrease in RTOP, RTAP and RTPP values indicates neuronal density reduction, and possibly axonal swelling (Le et al., 2020). These findings were anticipated, as we expected these tracts to deteriorate with aging. This validation underscored the neuroanatomical relevance of the connectome features identified by the FAGNN model as markers of brain aging and emphasized white matter integrity as a key factor in aging and neurodegeneration.

**Figure 7.**
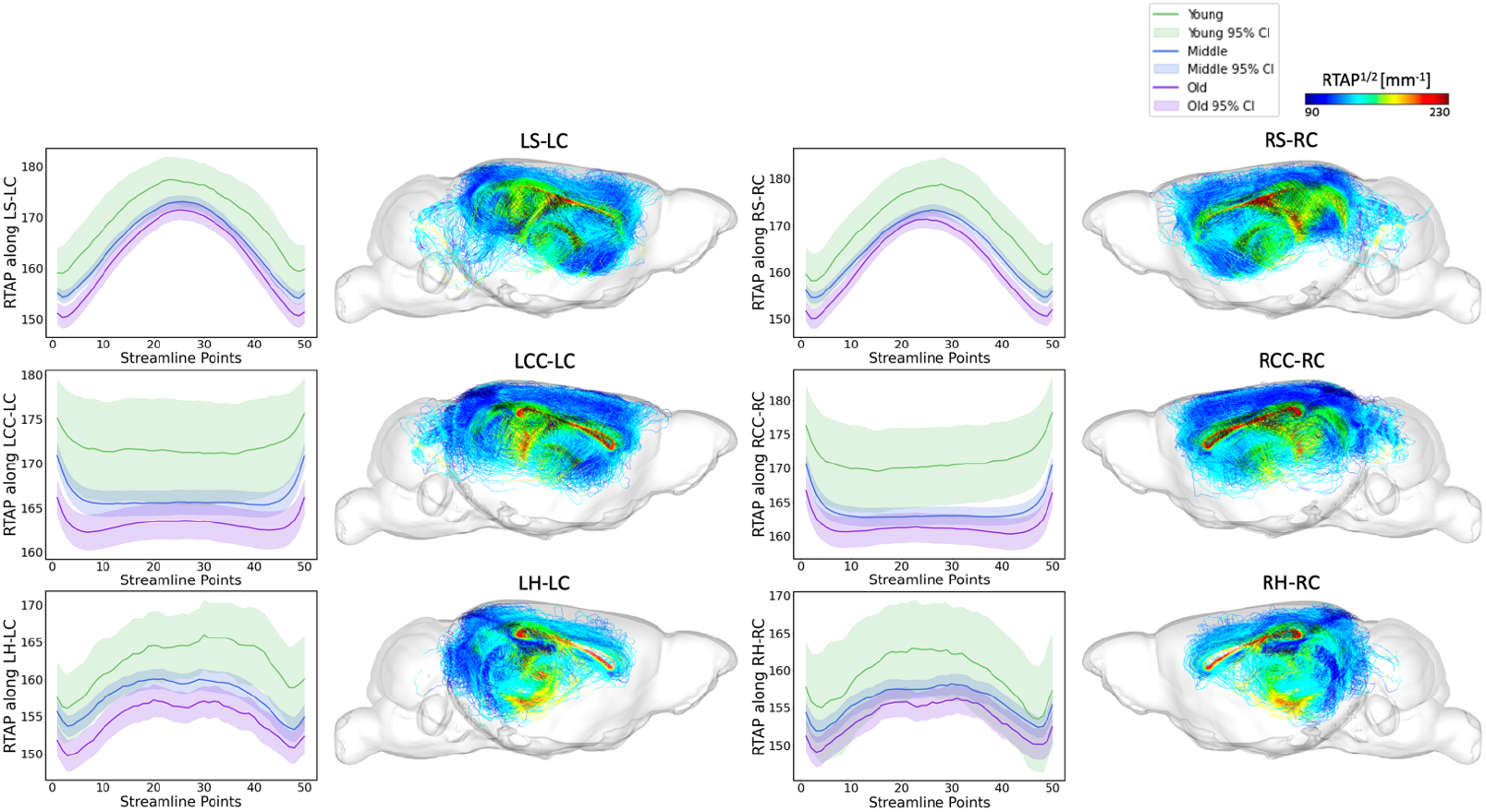
RTAP along tract profiles for the top 6 edges identified by FAGNN. The first row shows striatum-cingulum tract, second row shows corpus callosum-cingulum tract and last row shows hippocampus-cingulum tract. First column shows RTAP profile of left-left connection, second column shows the corresponding tractography with RTAP values. Third column shows RTAP profile of right-right connection, and the last column shows the corresponding tractography with RTAP values. The RTAP values along each tract for age groups were all significantly different with p<0.001.

**Figure 8.**
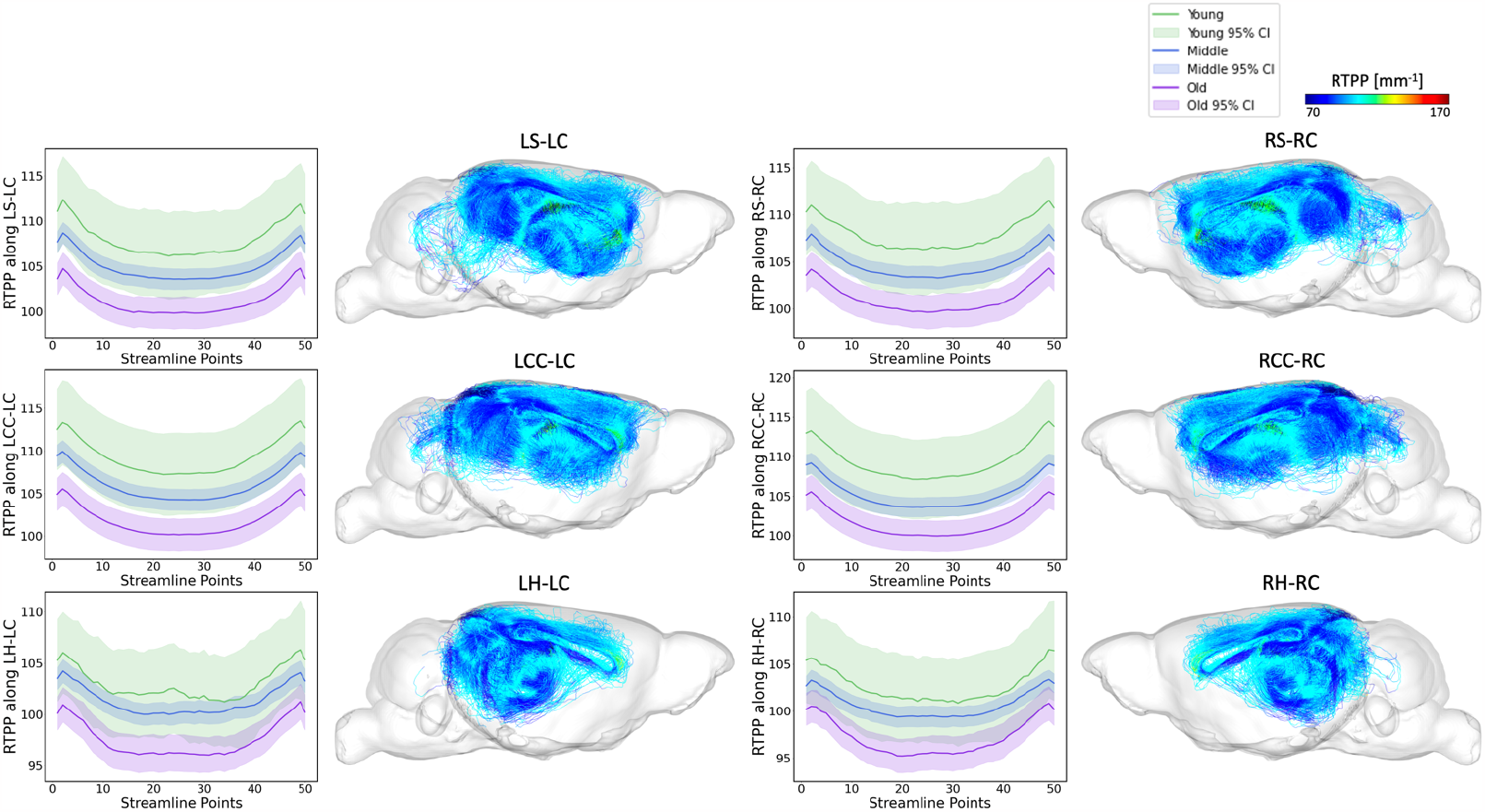
RTPP along tract profiles for the top 6 edges identified by FAGNN. The first row shows striatum-cingulum tract, second row shows corpus callosum-cingulum tract and last row shows hippocampus-cingulum tract. First column shows RTPP profile of left-left connection, second column shows the corresponding tractography with RTPP values. Third column shows RTPP profile of right-right connection, and the last column shows the corresponding tractography with RTPP values. The RTPP values along each tract for age groups were all significantly different with p<0.001.

The potential impact of our study consists in showing proof-of-principle that integrating the FAGNN model with clinical data may advance the early detection of AD, enhance the precision of disease staging, and tailor intervention strategies more effectively. The FAGNN model could prove helpful for tracking the progression of brain-related diseases and assessing the effectiveness of treatments over time. Moreover, the specific tracts identified in this study that connect the cingulum to other important regions require further validation through larger studies. Future studies that enhance the integration of various modalities could significantly contribute to the progress of AD research.

Nonetheless, our study is not without limitations. Transitioning from rodent models to human applications is an important but difficult procedure (Casey et al., 2015; Pappas & Nagy, 2019), that would benefit from cross-species studies. The results of this study could be tested through longitudinal human studies for subjects with different AD risk factors. Our results suggest that expanding the model with more comprehensive data, e.g. from omics studies, can improve prediction accuracy and robustness, thereby yielding more reliable results (Mahood et al., 2020; Miotto et al., 2017), and refining predictive accuracy. We need to consider the higher heterogeneity of human subjects that may contribute to brain aging, and that using large public databases on human aging such as UK Biobank (Sudlow et al., 2015) can help refine models, while mouse data is scarce. We recognize that the sample size of our current study is modest, especially in comparison to human datasets, but the genetic heterogeneity is also reduced and the environment in which these mice were raised was controlled. Our model’s inclusion of multiple datasets and the consequent increase in the number of parameters could complicate the model training. Also, as we included four risk factors, as well as continuous age for prediction, there are thus numerous permutations of various risk factor data, and the distribution of age range of mice for each group is not perfectly balanced. Despite these challenges, our findings demonstrate that the integration of risk factors and behavioral data can enhance prediction accuracy and precision, even as the model grows in size and complexity.

In conclusion, our research presents a comprehensive model for predicting brain age, incorporating a diverse set of data, from structural connectomes to environmental, genetic, and cognitive factors. We have revealed sub-networks that correlate to brain aging, and that multivariate data integration is beneficial for predictive modeling. These efforts represent a significant stride toward more personalized medicine in managing neurodegenerative diseases, and pave the way for future studies aimed at validating and broadening the application of connectomic biomarkers for aging and age-associated neurodegenerative diseases, such as AD.

## Data and Code Availability

The raw connectome, behavioral data and associated metadata are available at https://zenodo.org/records/10372075. The code necessary to reproduce the original analyses is available from https://github.com/hsmoon0/FAGNN.

## Author Contributions

Statistical and computational analysis was conducted by HSM, AM, JAS, RJA; primary writing and editing by HSM, AB; supervision by AB; and funding acquisition by CTB, AB. All authors have read and approved the final manuscript.

## Acknowledgment

We acknowledge support from the research staff in the Radiology department: Gary Cofer and Wyatt Austin, and BIAC staff: Chris Petty and Francis Favorini.

